# Systemic Analysis of Patient Derived Esophageal Epithelial Organoid Model Reveals Cell State Heterogeneity and Conservation of Differentiation Hierarchy

**DOI:** 10.64898/2026.10.29.762343

**Authors:** Alyssa Baccarella, Yusen Zhou, Tatiana A. Karakasheva, Ravi Gautam, Megha Lal, Jarad Beers, Zoe Mrozek, Ann Semaeo, Leena Mirchandani, Ryugo Teranishi, Takefumi Itami, Kanak V. Kennedy, Mark Mahon, Ezuka Ogbolu, Judith R. Kelsen, Kelly A. Whelan, Joshua B. Wechsler, Amanda B. Muir, Melanie A. Ruffner

**Author notes:** Corresponding Author Information: Melanie Ruffner, Address: Children’s Hospital of Philadelphia 3615 Civic Center Blvd, ARC 1202C Philadelphia, PA, 19104.

## Abstract

**Background and Aims:** Eosinophilic esophagitis (EoE) features prominent epithelial dysregulation that persists despite resolution of eosinophilia. Interleukin-13 (IL-13) is known to be a key upstream driver of epithelial abnormalities in EoE. We sought to benchmark an IL-13 esophageal organoid model against the *in vivo* epithelial state in EoE at a single cell resolution, to define in the role of IL-13 in aberrant epithelial development and inflammation in EoE, and to validate the IL-13 organoid as a model system to study the epithelial changes seen in EoE.

**Methods:** We performed scRNA-Seq on esophageal biopsies from control and EoE patients, and esophageal biopsy-derived organoids with or without IL-13. We compared cellular composition, differentiation trajectories, pathway enrichment, and differential gene expression within both systems, and assessed findings at a protein level.

**Results:** Seven distinct epithelial cell clusters were identified and preserved in both biopsies and organoids. EoE biopsies demonstrated expansion of cycling cells, whereas IL-13–treated organoids showed shift in the composition of the suprabasal compartments. Pseudotime topology was preserved across tissues, with a shift away from terminal differentiation in both EoE biopsies and IL-13–treated organoids. IL-13 reproduced changes in the transcription of central EoE genes and the dysregulation of epithelial proliferation and differentiation pathways seen in EoE. There was broad activation of immune related pathways within both EoE biopsies and IL-13-treated organoids, however, biopsies additionally showed activation in several immune and extracellular matrix remodeling programs that IL-13 alone did not reproduce.

**Conclusion:** IL-13–treated esophageal organoids replicate key epithelial features of EoE including dysregulated epithelial development, impaired cornification, and inflammatory signals, while preserving lineage architecture, providing a tractable platform for epithelial-targeted mechanistic and therapeutic studies.

**Background:** IL-13-treated esophageal organoids provide a model for studying epithelial dysfunction in eosinophilic esophagitis (EoE), but their ability to reproduce disease-associated changes at single-cell resolution remains unclear.

**Findings:** Esophageal organoids preserved epithelial differentiation trajectories, while IL-13 treatment recapitulated key EoE features, including impaired terminal differentiation, retention of basal-like programs, and activation of inflammatory pathways.

**Limitations:** Organoids contained fewer quiescent basal and terminally differentiated cells than biopsies and did not fully reproduce the broader inflammatory and tissue-remodeling programs observed in EoE.

**Impact or Future Directions:** These findings establish esophageal organoids as a model for investigating epithelial dysfunction in EoE, providing a foundation for mechanistic studies, targeted perturbations, and future therapeutic testing.

## INTRODUCTION

Eosinophilic esophagitis (EoE) is often characterized by a mixed inflammatory infiltrate ^1,2^ and epithelial abnormalities including basal cell hyperplasia and dilated intercellular spaces that impair barrier function allowing for ongoing antigen exposure^3,4^. These epithelial changes persist despite resolution of eosinophilia and are associated with ongoing symptoms and mast cell infiltrate^5,6^. The majority of transcriptional changes in the EoE transcriptome arise from the dysregulated epithelium,^7^ and studies have previously demonstrated heterogeneity within the esophageal epithelium in homeostasis and disease^8–10^. Thus, understanding the molecular changes in the epithelium is critical to understanding disease pathophysiology and identifying targets to restore homeostasis.

Interleukin-13 (IL-13), a key type 2 cytokine in EoE^11^, promotes basal cell hyperplasia, impaired epithelial differentiation, and barrier disruption^12–14^. Esophageal organoids are a powerful tool to investigate IL-13-mediated epithelial responses and ultimately test therapeutic targets to re-establish epithelial homeostasis^9,15,1619^. However, the extent to which this model recapitulates the *in vivo* esophageal epithelium has not been systematically assessed at a single-cell resolution.

Here, we compare single-cell transcriptomic profiles of control and EoE esophageal biopsies with untreated and IL-13-treated patient-derived organoids. We assess cellular composition, differentiation trajectories, and transcriptional programs to define the features of disease driven by IL-13 and identify limitations of the organoid model.

## MATERIALS AND METHODS

Patients and Esophageal Biopsies: This study received approval from the institutional review boards at the Children’s Hospital of Philadelphia and Lurie Children’s Hospital of Chicago.

Informed consent was obtained before esophagogastroduodenoscopy. EoE was defined by symptoms of esophageal dysfunction and ≥15 eosinophils per high-powered field (eos/hpf)^2^. Patients in histologic remission were excluded. Controls had no history of EoE or other gastrointestinal disease and normal histopathology. Biopsies were collected and transported in RPMI with 10% fetal bovine serum and dissociated as described previously^17^.

Organoid Cultures: Organoids from human esophageal biopsies were established as previously described^15^. Dissociated cells were embedded in Matrigel (Corning) and keratinocyte serum free medium (KSFM) with 1.8 mM Ca^2+^ (KSMFC), then incubated at 37° for 30 minutes. On day 1, solidified wells received KSFM with 10 µM Y-27632 (ROCK inhibitor; Sigma-Aldrich); medium without Y-27632 was added every other day. From day 7, organoids were cultured with or without IL-13 (10 ng/mL) ^9^ and harvested on day 11.

scRNA-Seq library preparation: Single-cell suspensions were prepared from biopsies and organoids. Biopsies were washed three times in RPMI with 10% FBS, then washed with ethylenediaminetetraacetic acid (EDTA) stripping buffer (5 mM EDTA, 10mM 4-(2-hydroxyethyl)piperazine-1-ethane-sulfonic acid (HEPES), 10 mM dithiothreitol (DTT), 10% FBS in calcium-free Hank’s Balanced Salt Solution (HBSS), then washed 3 times with HBSS with HEPES. Tissue was minced and digested with collagenase IV and DNAse and filtered. If needed, dead cells were removed (Dead Cell Removal Kit, Miltenyi Biotec). Single-cell cDNA libraries were generated using Single Cell 5’ Reagent Kits v1 (10x Genomics) per manufacturer instructions and sequenced on an Illumina Novaseq6000 sequencer.

Organoids were harvested in dispase (1:5), disrupted by vigorous pipetting, centrifuged, and washed. Pellets were then incubated with trypsin to generate single-cell suspensions, followed by quenching with soybean trypsin inhibitor (STI), centrifugation, and resuspension in culture medium. If needed, dead cells were removed. Single-cell cDNA libraries were generated using Single Cell 3’ Reagent Kits v2 (10x Genomics) per manufacturer instructions and sequenced on an Illumina NovaSeq6000.

Single-cell RNA-Sequencing (scRNA-seq) Data Processing Data was processed in CellRanger 3.1.0 and Seurat v4 (see Supplemental Methods). Only epithelial cells were retained for downstream analysis, based on known marker genes. Cluster labels were assigned based known cell markers curated from the literature as well as examination of marker genes, identified by FindClusterMarkers^17,18^. A resolution of 0.6 was used for clustering, and several clusters were combined based on the known recognized epithelial cell types^17,18^. Following cluster assignment and labeling, the active EoE and IL-13-treated organoid samples were normalized using SCTransform normalization and then projected onto the control reference trajectory using FindTransferAnchors and MapQuery (Seurat)^19^.

Quantification of cellular populations: Frequencies of cluster populations were tabulated per patient. The Kruskal-Wallis test was used to compare the medians of individual groups. *P*-value less than 0.05 was considered significant.

Sub-clustering and pathways analysis of differentiated cell population: UMAP visualization revealed distinct subpopulations of cells with sufficient cell numbers for subclustering analysis of the Diff2 group in the Control subject group. However, this was not seen in other conditions, where the number of cells in Diff2 was markedly reduced. To perform subclustering of the Control Diff2 population, the Diff2 cells were subdivided using Seurat’s FindSubCluster() function at a resolution of 0.1. Marker genes were identified using one-vs-rest comparisons to define transcriptional programs uniquely enriched within each subpopulation using FindAllMarkers(). Significant genes (p-adjusted < 0.05) were subsequently subjected to over-representation analysis with the enrichGO() function from ClusterProfiler^20^, using default parameters. Gene Ontology (GO) biological pathways were retrieved from MSigDB v2025.1.Hs^21,22^, and pathways were considered significant at an adjusted p-value < 0.05.

Pseudotime Analysis: Monocle 3 (version 1.0.0)^23^ was used to perform trajectory inference analysis on the integrated scRNA-seq data. Based on prior knowledge, we set the transit-amplifying (TA) cluster as the root to estimate pseudotime trajectories^10^.

Basal and Differentiation Module Scores: Genes defining basal or differentiated scores were determined using FindMarkers() to compare the quiescent basal and diff 2 clusters to all other clusters in the control esophagus biopsies. These genes (Supplemental Table 1) were then used to compute module scores using AddModuleScore() within Seurat.

Cell Cycle Prediction: A cell cycle score was assigned to each cell based on the expression of G2/M and S phase markers using the CellCycleScoring() function in Seurat^24^.

Differentially Expressed Gene Analysis: Raw gene expression counts were aggregated by cell type within each sample. The reorganized gene expression counts were normalized, and differentially expressed genes (DEGs) were identified using the DESeq2 package (version 1.44.0)^25^. Benjamini-Hochberg adjusted *p-*value less than 0.05 was considered significant. The stat output field from DESeq2 was then used as input for GSEA pre-ranked analysis to identify enriched pathways from the GO biological processes databases^26^, with q-values less than 0.05 considered significant. Pathways were compared for shared significance between tissues at the individual pathway level, also assessing for similarity with closely related pathways, defined as Jaccard index ≥ 0.5. Contributing leading edge genes were assessed. Pathways were manually curated utilizing pattern-matching on term names into families for visualization (Supplemental Table 2). Families were classified as shared, biopsy or organoid dominant (at least 1 family pathway significant in ≥ 3 clusters in one tissue and ≤1 in the other tissue) or biopsy/organoid only.

For leading-edge visualizations, a gene was considered "present" if it appeared in the leading edge of any pathway in a specific tissue in at least one cluster where that pathway was significantly enriched.

Transcription Factor Analysis: Transcription factor (TF) enrichment analysis was performed using the ChEA 2016 database via the Enrichr R package^27^. Shared differentially expressed genes (DEGs) between biopsy and organoid samples were defined for each epithelial cluster based on pairwise intersections of significant genes (adjusted *p* < 0.05). These shared gene sets were queried using enrichr() with the ChEA 2016 database, including results only from human datasets. Significantly enriched TFs were defined based on adjusted p < 0.05.

Immunohistochemistry and Immunofluorescence: Organoids were fixed and embedded as previously described^9,15^ then subjected to immunohistochemistry and immunofluorescence.

Staining was done on the Leica BOND RX^m^ Automatic slide stainer. For immunohistochemistry, a heat-induced epitope retrieval, pH 6, with citric acid was used. Slides were stained with primary anti-cornulin antibody (1:1500, rabbit polyclonal, ab224122, Abcam), anti-Ki-67 antibody (1:100, rabbit monoclonal, ab16667, Abcam), anti-TP63 antibody (1:1000, ab124762; rabbit monoclonal Abcam), or anti-SOX2 antibody (1:100, rabbit monoclonal, 14962S, Cell Signaling Technology), and antigen detection was done with the Bond Polymer Refine Detection Kit (Leica Biosystems).

For immunofluorescence, sections were stained with primary anti-IVL monoclonal antibody (1:100, I9018; mouse monoclonal, Millipore Sigma) and nuclei were stained with DAPI, then with Cy3 AffiniPure Donkey Anti-Mouse IgG secondary antibody (1:500, AB_2349813, Jackson ImmunoResearch).

Multiple 40x HPF images were taken with the M7000 Evos Imaging System (ThermoFischer Scientific) and processed uniformly with Evos Analysis software (ThermoFischer Scientific) and ImageJ. ImageJ was utilized to quantify the percent relative area positivity (cornulin) or percent positive nuclei (SOX2, MKI67, TP63)^28^ and Mann Whitney Test was used to compare, with *P*-value less than 0.05 considered significant.

Data Availability: Esophageal biopsy data will be deposited to dbGAP and accessible upon request or via a data use agreement. Organoid data is deposited in the Gene Expression Omnibus under GSE227593.

## RESULTS

### Patient-Derived Esophageal Organoids Contain All Major Epithelial Cell Types

We generated three esophageal epithelial organoid cultures from independent healthy control donors. Each organoid culture was grown under baseline conditions or stimulated with IL-13 prior to performing single cell RNA-Sequencing (scRNA-seq)^15^. Whole biopsy scRNA-seq was also performed on 5 control patients and 14 patients with active eosinophilic esophagitis (EoE) (Figure 1). In this cohort, we observed a higher proportion of males in the EoE group, consistent with prior studies (Table 1). There were no differences in atopic diseases (Table 1). Most patients were on treatment, including diet elimination and proton pump inhibitor (Supplemental Table 3). Additional endoscopic and histologic findings are displayed in Supplemental Table 3.

**Figure 1.**
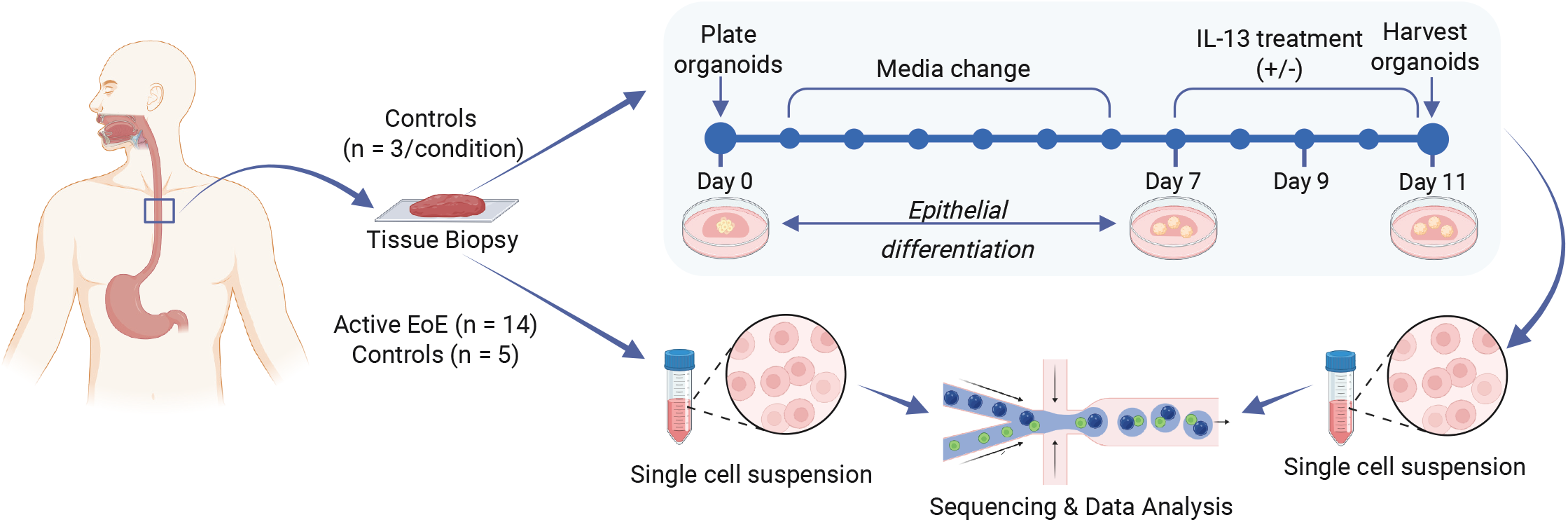
Experimental Design. Schematic of experimental design.

**Table 1:**
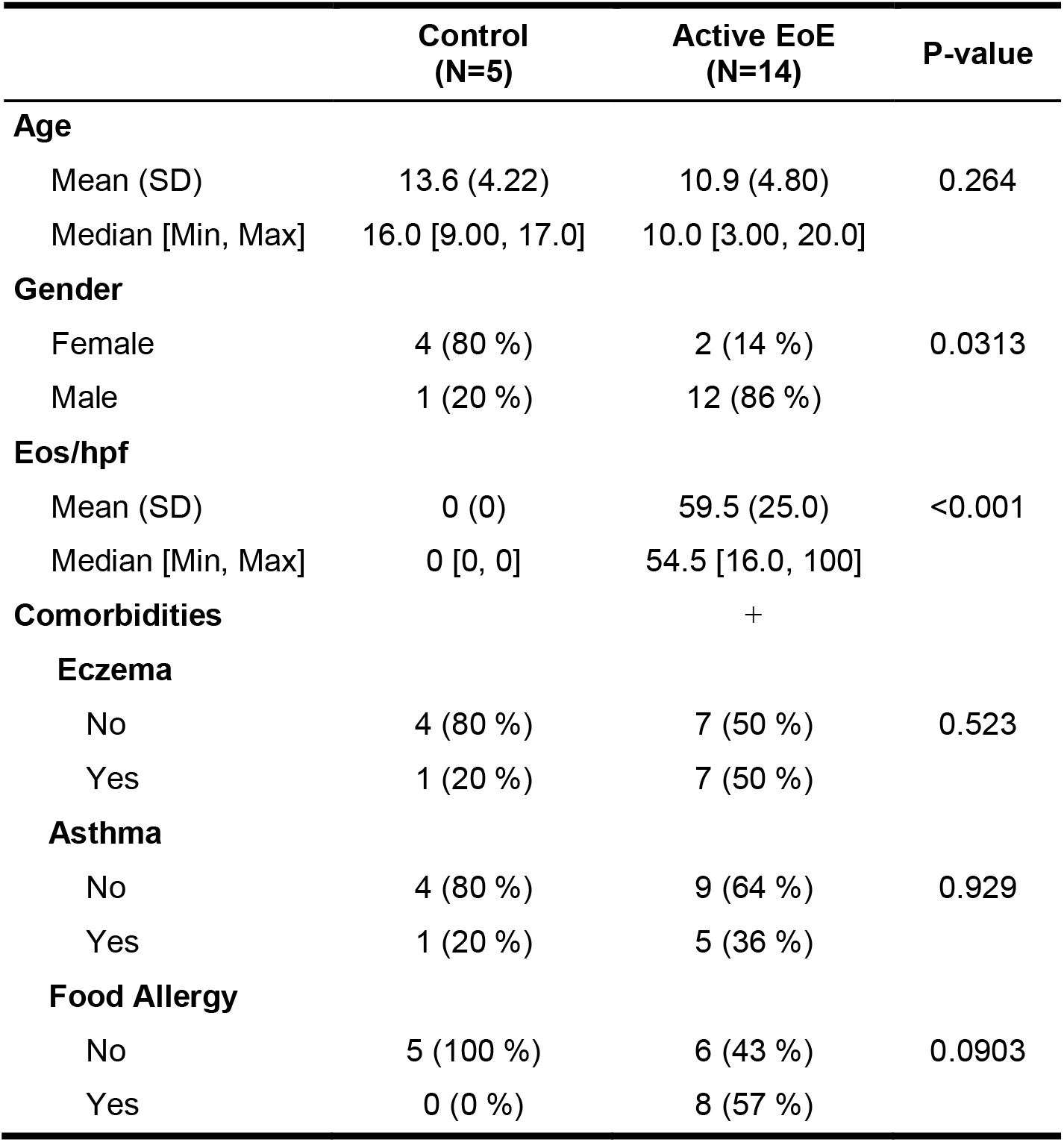
Patient Characteristics Biopsy Data.

After quality control filtering, a total of 25,538 epithelial cell transcriptomes from biopsies and 41,157 cell transcriptomes from organoids were retained for downstream analysis (Supplemental Figure 1) and seven distinct clusters were identified. Clusters were visualized using UMAP plots (Figure 2A) and assigned identities based on both canonical markers^9,17,18^ and specific enrichment within the cluster in non-EoE controls (Figure 2B, Supplemental Table 4). We identified five early clusters (sharing *DST, COL17A1,* and *KRT15)* including two basal clusters (*IL1R2*, *TSLP*): quiescent basal (*PDPN, CXCL14*) and slow cycling basal (emergence of low level *CCNB1, TOP2A, MKI67)*; a transit-amplifying (TA) cluster (high expression of *MIK67, TOP2A, HMGB2, and RRM2)*, and two suprabasal clusters sharing *DSC2, SERPINB3,* with the emergence of *HES2* and persistence of *KRT15*, delineated by decreased *CNNB1* and the emergence of *C10orf99, DEFB1,* and *IVL* in suprabasal 2. We also detected two differentiated clusters, which no longer expressed proliferation-related genes and instead showed expression of markers such as *EVPL and IVL,* and ultimately markers of terminal differentiation *(CRNN, FLG)*.

**Figure 2.**
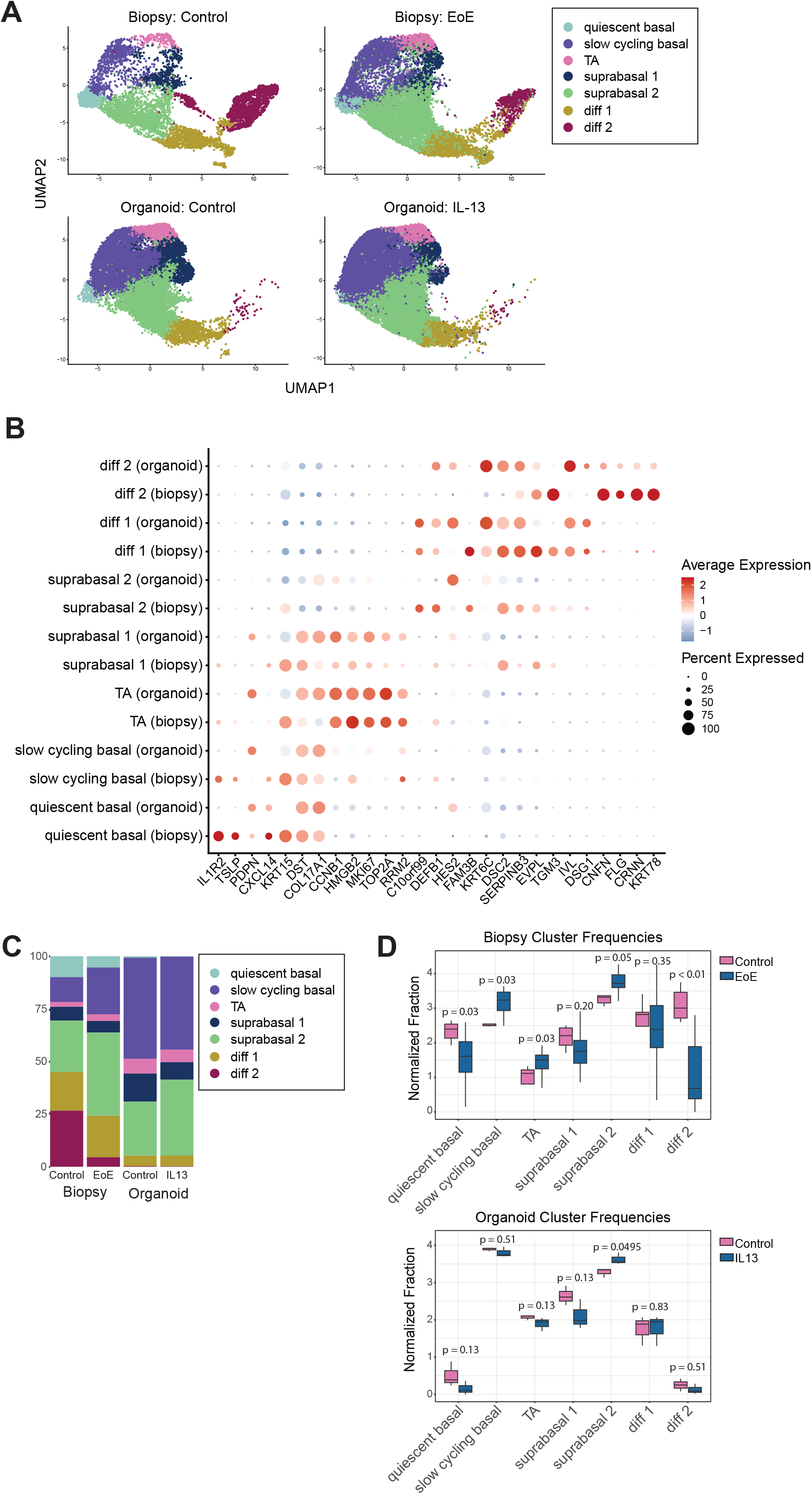
Esophageal organoids recapitulate the cell populations seen within healthy esophageal epithelium, with expansion of cycling cells. A. UMAP plots of single-cell gene expression data from biopsies and organoid data B. Marker genes defining each cell population in control biopsies and control organoids C. Relative cluster frequencies by condition D. Relative ln(fraction + 1) cluster frequencies by disease/treatment condition in biopsy (left) or organoid (right), *p-*values shown.

All epithelial clusters identified in biopsies were also detected in organoids under both control and IL-13 treatment conditions. However, organoids demonstrated a relative reduction in quiescent basal and differentiated cell populations compared to biopsies (Figure 2C,D; Supplemental Figure 2), likely reflecting limitations of the *in vitro* system. Consistent with this, organoids exhibited significantly increased proportions of dividing cells (slow-cycling basal and TA cells; p < 0.001) and decreased quiescent basal (p < 0.001) and differentiated cells (p = 0.011) relative to biopsies (Figure 2C; Supplemental Figure 2).

We next compared cluster frequencies across disease conditions within each specimen type. In biopsies, EoE samples showed an increase in cycling populations (slow-cycling basal and TA clusters) and a decrease in quiescent basal cells compared to controls. There was also a trend toward more suprabasal 2 cells (p = 0.05). Notably, EoE biopsies demonstrated a significant reduction in the late differentiated Diff2 cluster (p < 0.01), consistent with prior reports of impaired terminal differentiation in EoE^17,29^. In organoids, by contrast, only suprabasal 2 cells were significantly increased following IL-13 treatment compared to control (Figure 2D).

Given the pronounced reduction in Diff2 cells observed in EoE biopsies (Figure 2D; Supplemental Fig. 3A), we sought to further characterize Diff2. We focused subsequent analyses on biopsy samples, as organoids contained very few late differentiated cells, precluding meaningful sub-clustering. UMAP visualization of control biopsies suggested distinct subpopulations within Diff2 that appeared selectively depleted in EoE. Sub-clustering identified four populations within Diff2; although one small cluster (n = 2 cells) was excluded from further analysis. Of the three remaining subclusters, two (Diff2_0 and Diff2_2) were preferentially reduced in EoE compared to control (Supplemental Figure 3B).

The preserved subcluster (Diff2_1) was characterized by elevated expression of keratinization-associated genes, including *FLG*, as well as genes involved in RNA splicing, ATP synthesis, and protein localization (Supplemental Figure 3C). One depleted population (Diff2_0) exhibited relatively few distinguishing markers (Supplemental Table 4), though enriched genes were associated with immune responses (*S100A8*, *S100A9*) and cornification (small proline-rich protein family genes). The second (Diff2_2) showed enrichment of ribosomal function, regulation of peptidase activity, detoxification pathways, and response to metal ions (Supplemental Figure 3C), consistent with a previously described metallothionein-rich differentiated epithelial population reported to be reduced in EoE^17^.

Together, these findings indicate selective depletion of specific terminally differentiated epithelial subsets in EoE, highlighting potential functional specialization within the Diff2 compartment and suggesting avenues for further mechanistic investigation.

### Biopsy Epithelium and Organoids Show Similar and Broad Expression of IL-13 Receptor

Given the use of IL-13 treatment to model EoE-associated epithelial changes, we next sought to determine which epithelial populations have the capacity to respond to IL-13.

Expression of IL-13 receptor components *IL13RA* and *IL4R* was broadly detected across epithelial clusters in both biopsy and organoid conditions, although *IL4R* expression was lower in terminally differentiated cells (Supplemental Figure 4). Neither gene demonstrated significant differential expression between control and EoE biopsies or between untreated and IL-13-treated organoids within any epithelial cluster (Supplemental Table 6). These findings suggest that the capacity for IL-13 signaling is broadly distributed across esophageal epithelial populations and is not restricted to a specific differentiation state.

### EoE Biopsies and IL-13-Treated Organoids Show Stalled Differentiation Trajectories

We next performed pseudotime analysis to infer the relative differentiation states of cells under each condition. We set the TA cluster set as the starting node, reflecting the capacity of TA cells to either progress through differentiation or return to a more stem-like phenotype under stress^16^. The analysis revealed several distinct trajectories originating from the cycling populations. One branch terminated in the quiescent population, while another represented progressive differentiation through the suprabasal populations and into the terminally differentiated states. In addition, a shorter but distinct trajectory connected the TA and slow-cycling clusters (Figure 3A).

**Figure 3:**
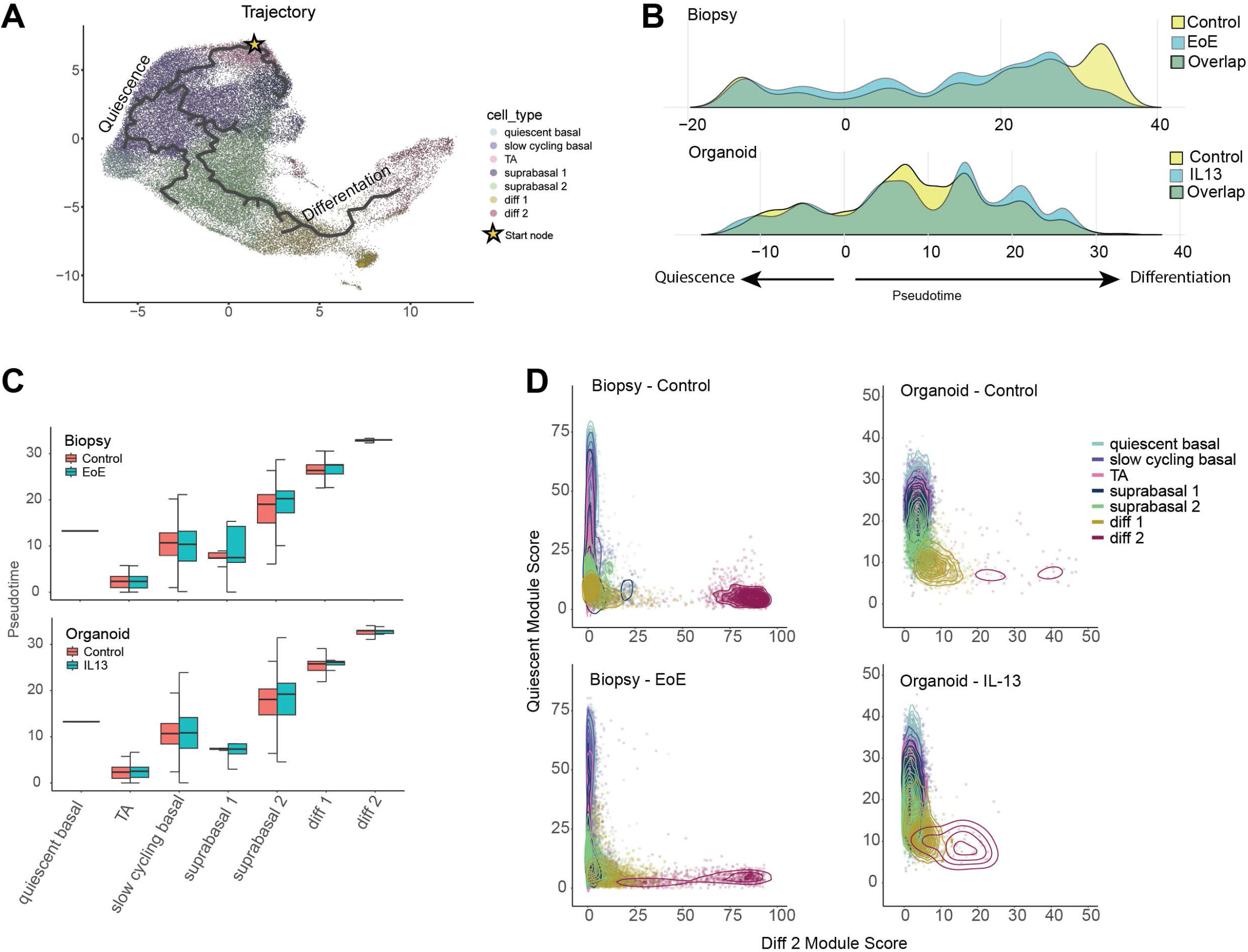
EoE esophagus and IL-13-treated organoid show preservation of differentiation trajectory with aberrant terminal differentiation. A. Monocle 3 predicted pseudotime trajectories overlayed on integrated UMAP, star indicates root node B. Frequencies of cells at a given pseudotime (y axis) in either biopsies or organoids C. Box plots of pseudotime values by cluster D. Density plots of quiescent and differentiated module scores, by cluster, by tissue and condition.

We examined the density distributions of cells from each condition along the pseudotime trajectory. Compared with biopsies, organoids had higher cell density earlier in the pseudotime axes and fewer cells at the termini of both branches, consistent with a greater proportion of proliferative and less differentiated cells in organoids at day 11 (Figure 3B). There were no notable differences in untreated and IL-13-treated organoid cell densities. Within biopsies, there was a decrease in cell density along the most terminal differentiation pseudotime axis in EoE biopsies compared to control biopsies (Figure 3B).

To distinguish changes in the abundance of differentiation states from alterations in the differentiation trajectory itself, we then examined the distribution of pseudotime within each cluster. As expected, more differentiated populations exhibited higher pseudotime values, and this relationship was preserved across all biopsy and organoid conditions (Figure 3C). There were no significant differences in the distribution of pseudotime values within a given cluster when comparing EoE biopsies or IL-13-treated organoids to controls (Figure 3C). Thus, the reduced density of cells at the terminal end of pseudotime primarily reflects a reduction in the abundance of terminally differentiated cells, particularly the diff2 population, rather than a shift in the sequence of individual differentiation states.

Together, these findings show that although organoids contain a higher proportion of proliferating cells and fewer terminally differentiated cells than biopsies, cells in both systems progress through similar transcriptional programs, as demonstrated by preserved pseudotime trajectories across conditions. These results support the ability of the organoid model to recapitulate the biologic differentiation trajectory of the esophageal epithelium.

While the relative differentiation trajectories are preserved in organoids compared to the esophageal epithelium, it has previously been reported that in EoE, suprabasal cells and terminally differentiated cells maintain progenitor identity programs, which may contribute to the epithelial derangements seen in EoE^10^. To further investigate epithelial differentiation, we developed gene modules preferentially enriched in either quiescent basal or terminally differentiated cells from non-EoE control patient biopsies (Supplemental Table 1). Contour plots mapping the quiescent module score versus the differentiated module score revealed a notable decrease in differentiation scores within the differentiated populations in both EoE biopsies and IL-13-treated organoids. Furthermore, the diff 1 population shifted towards a higher quiescence score in IL-13-treated organoids (Figure 3D). This suggests a decrease in terminal differentiation programs in EoE, which is recapitulated in the IL-13 organoid model.

### EoE Biopsies and IL-13-treated Organoids Show Similar Dysregulation of Proliferation and Differentiation

We next examined gene expression patterns within the cell populations to identify potential shared patterns of epithelial disruption that lead to a less differentiated state in EoE and the IL-13-treated organoid model. Gene set enrichment analysis (GSEA) of Gene Ontology (GO) biological processes was performed by cluster, comparing EoE biopsies or IL-13 organoids with their respective controls (Figure 4A, Supplemental Table 7). Overall, biopsies across all clusters showed many more significant GO terms as compared to organoids (4860 versus 822). Two organoid clusters, quiescent basal and diff 2, yielded fewer significant terms (29 and 14), likely due to the lower cell recovery from these clusters. Because GO-BP terms are extensively nested, we next assessed specificity at three levels: the individual pathway level, closely-related terms, and curated pathway families (Supplemental Table 8).

**Figure 4.**
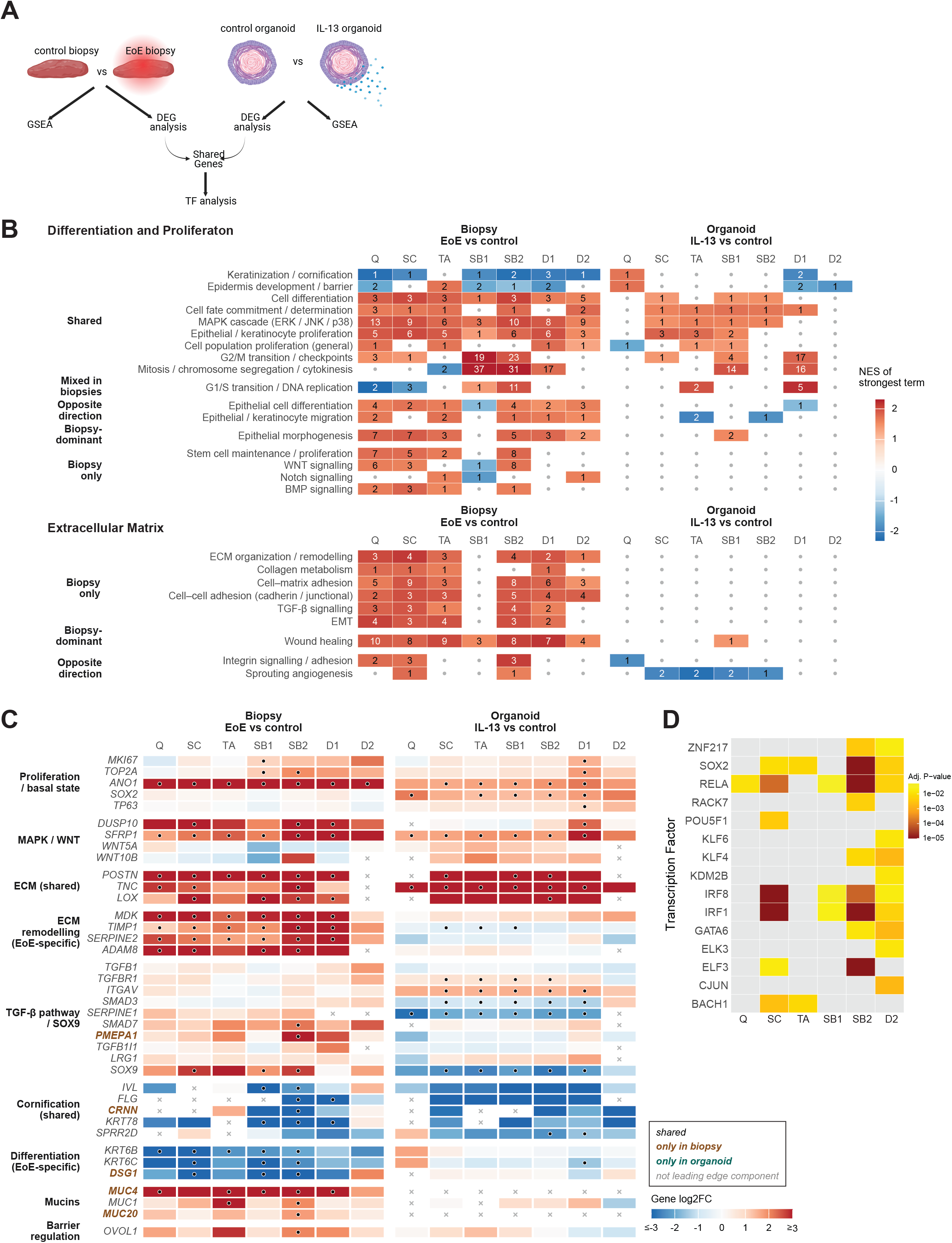
IL-13-treated organoid recapitulates dysregulation of proliferative pathways and basal cell transcription factor expression seen in EoE. A. Analysis schematic B. Epithelial proliferation, differentiation, and extracellular matrix related pathway families are significantly enriched in both EoE biopsies and IL-13-treated organoids. Pathways were collapsed into families by theme, and the strongest NES of a pathway-family is shown. The number indicates the number of pathways significant in that family (q <0.05) C. Select leading edge genes of significant pathways. Color indicates log2 fold change. Circle = p adjusted <0.05. D. TF analysis (ChEA 2016) of DEGs shared in both organoids and biopsies by cluster.

Pathways related to epithelial cell proliferation were dysregulated in both EoE biopsies and IL-13-treated organoids (Figure 4B). These included epithelial cell proliferation, cell fate commitment, and cell-cycle pathways such as G2/M transition and mitosis. The IL-13-inducible chloride channel, *ANO1*, which promotes basal cell proliferation in EoE^30^ was significantly upregulated across clusters in both EoE and IL-13-treated organoids (Figure 4C). MAPK signaling, which has been implicated in EoE pathophysiology^31^, was enriched in both biopsies and organoids. WNT signaling pathways only reached significance in biopsy, however, the WNT antagonist, *SFRP1,* recently shown to mediate aberrant proliferation in the EoE epithelium in a STAT3 dependent manner^12^ was induced in both EoE and IL-13 treated organoids (Figure 4C).

To identify candidate regulators of the shared differentiation defects, we compared differential expression between EoE and control biopsies and between IL-13-treated and control organoids. Differentially expressed genes were broadly concordant between the two systems. The quiescent basal and diff 2 clusters showed the least concordance, likely driven by the low numbers of cells detected in these populations in organoids (Supplemental Table 6, Supplemental Figure 5). To identify potential upstream regulators of these shared DEGs, we identified candidate transcription factors (TFs) in the union of differentially expressed genes by cluster. SOX2, RELA, IRF1, and IRF8 emerged as top predicted TFs (Figure 4D). *SOX2* has recently been implicated in mediating the retention of basal programs in EoE^10^ and contributed to the leading edge of proliferation pathways in both biopsy and organoid. At a transcriptional level, *SOX2* only reached significance in IL-13 treated organoid (Figure 4C), however, it has previously been shown to be in be enriched in EoE biopsies at the protein level^9^. Thus we further examined its protein expression pattern in organoid and found that the percentage of SOX2+ cells increased in IL-13-treated organoids as compared to control (Figure 5A). We also examined the protein expression of TP63, a key basal cell transcription factor^32,33^, which has previously been shown to be increased in suprabasal and differentiated cells in EoE^10^. Similar to SOX2, IL-13-treated organoids also had a significantly increased percentage of TP63-positive cells, suggesting that dysregulation of cell proliferative pathways results in a higher proportion of cells maintaining basaloid features in IL-13-treated organoids, similar to the basal cell hyperplasia seen in EoE. Surprisingly, we did not detect a change in the number of Ki-67+ dividing cells in the IL-13-treated organoids at day 11 (Figure 5A). In EPC2-hTERT-derived organoids, Ki-67 showed a trend toward increased expression in IL-13-treated organoids compared to control at day 9 (p = 0.12, Supplemental Figure 6), suggesting an earlier time point may be more appropriate for studying proliferative differences in organoids. Taken together, these results indicate that IL-13-treated organoids maintain general differentiation trajectories and exhibit perturbed proliferation pathways similar to those observed in the esophageal epithelium during EoE. This may partly be caused by increased SOX2 and TP63 activity and could explain the basal cell hyperplasia characteristic of EoE.

**Figure 5:**
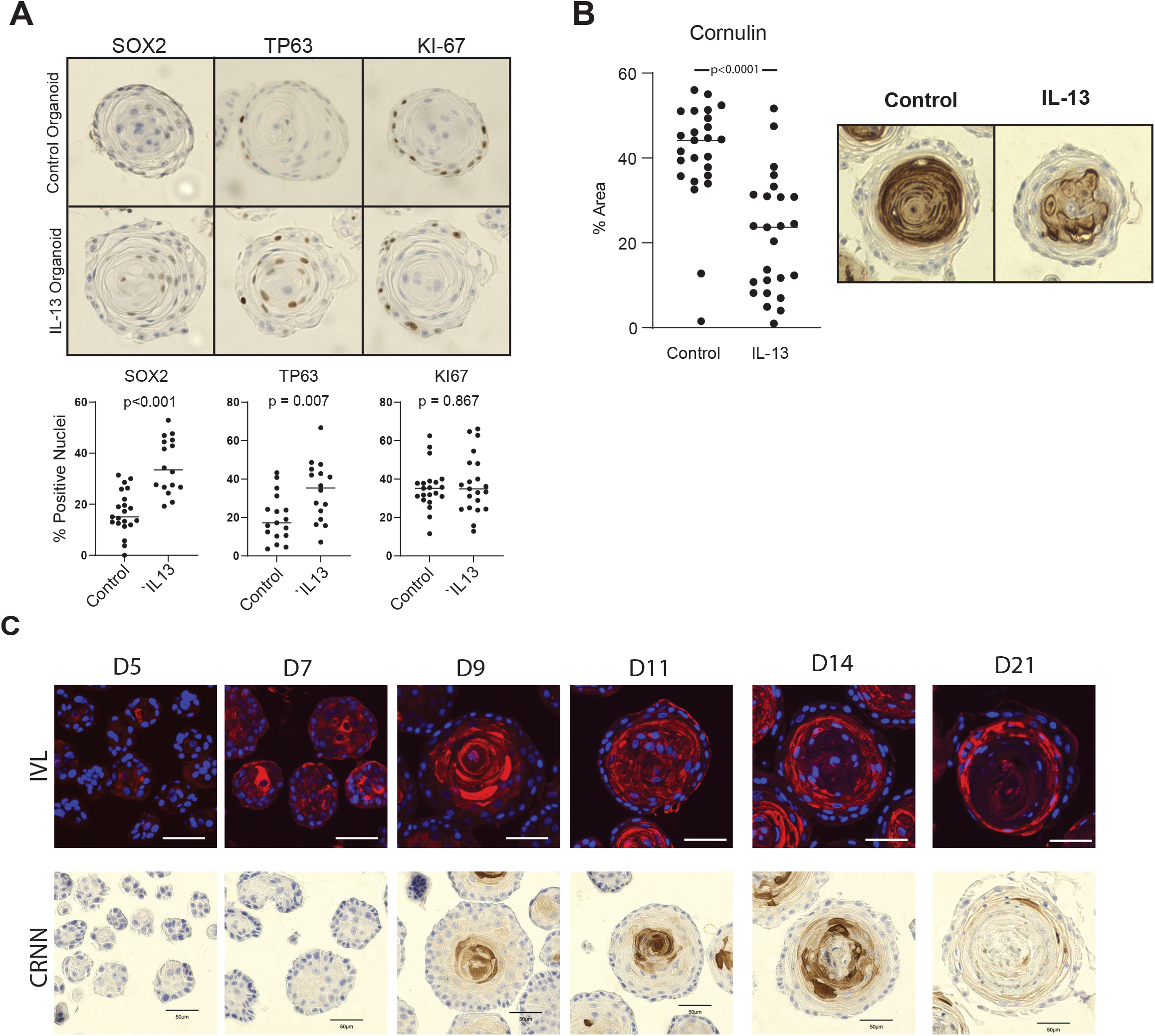
IL-13-treated organoids exhibit protein-level dysregulation of differentiation. A. Representative immunohistochemical staining for SOX2, TP63, and Ki-67 in organoids, quantified with *p*-values shown (Mann Whitney U). B. Quantification of cornulin staining in control vs IL-13-treated patient derived organoids, p-value shown (Mann Whitney U) C. Developmental time course of involucrin (red) and DAPI (blue) -top, and cornulin expression – bottom, in control EPC2 organoids. Bars represent 50um.

#### IL-13-treated Organoids Do Not Recapitulate Transforming growth factor beta (TGF)-β-associated remodeling programs

Because fibrosis is a known complication of EoE, we next investigated pathways involved in extracellular matrix (ECM) organization and remodeling. These pathways were nearly exclusively enriched in EoE biopsies but not in IL-13-treated organoids. ECM organization, collagen metabolism, cell matrix adhesion, TGF-β signaling, and EMT related gene sets were enriched across EoE biopsy clusters but were not enriched in IL-13-treated organoids (Figure 4B). IL-13 did, however, induce several extracellular matrix remodeling genes that were also induced in EoE, most notably periostin (*POSTN), TNC*, and *LOX* (Figure 4C). In contrast, other remodeling genes that contributed to the biopsy-specific pathways were induced only in EoE, including *MDK,* protease inhibitors *TIMP1* and *SERPINE2,* and metalloproteinase *ADAM8* (Figure 4C).

TGF-β is a central driver of fibrosis in EoE^34^. TGF-β-related pathways were enriched in EoE biopsies but were not enriched in IL-13-treated organoids. Although IL-13 treatment in organoid induced upregulation of TGF-β signaling machinery including *TGFB1R* and *ITGAV,* downstream transcriptional mediator, *SMAD3,* was downregulated, as was canonical TGF-β target *SERPINE1. SOX9,* which is induced by TGF-β and promotes fibrosis in other tissues ^35,36^, was also strongly induced in EoE but repressed in IL-13-treated organoids. Epithelial-mesenchymal transition (EMT) pathways were likewise only enriched in biopsies. Their leading edge included multiple TGF-β pathway genes (*TGFB1, TGFBR1, SMAD2/3, TGFB1I1*).

Together, these findings suggest that IL-13 alters esophageal epithelial cells’ capacity to respond to TGF-β, but is insufficient to induce TGF-β-associated remodeling or EMT, due to the lack of TGF-β from non-epithelial cells within the organoid microenvironment.

### IL-13-Treated Organoids and EoE Biopsies Show Disruption in Cornification

Disruption of the epithelial barrier and diminished terminally differentiated, cornified cells promote chronic antigen exposure and subsequent inflammation in EoE. Consistent with this, keratinization and cornification pathways were suppressed in both EoE biopsy and IL-13-treated organoids (Figure 4A). The leading edge genes in these pathways included multiple markers of terminal differentiation and cornification (Figure 4C), including the early differentiation marker *IVL* and genes associated with later stages of differentiation and keratinization, including filaggrin (*FLG)*^37^, keratins (*KRT6B, KRT6C,* and *KRT78*), and small proline-rich protein, (*SPRR2D*).

Membrane-bound mucins *MUC4, MUC1* and *MUC20* were induced in EoE biopsies.

Only *MUC1* contributed to pathway leading edges in organoids, but it did not reach significance in any cluster (Figure 4C). Mucin genes were generally expressed at low levels in the organoid model. Together, these findings demonstrate a shared pattern of impaired terminal differentiation and keratinization in EoE that is largely recapitulated in IL-13-treated organoids.

The transcription factor *OVOL1* was identified in the leading edge of the epidermal cell differentiation pathway. *OVOL1* was recently shown to positively regulate Serine peptidase inhibitor Kazal type 7 (*SPINK7),* which promotes barrier integrity and is depleted in EoE ^38^. IL-13 induces *CAPN14* and promotes OVOL1 degradation, leading to loss of SPINK7 and subsequent epithelial barrier disruption and increased TSLP production^39^. Consistent with this, *CAPN14* was significantly upregulated and *SPINK7* was downregulated in EoE biopsies, with *SPINK7* expression largely restricted to the differentiated clusters. By contrast, *TSLP* expression was highest in the basal populations, but it did not differ significantly between control and EoE biopsies. In organoids, *CAPN14* and *SPINK7* showed directional changes similar to those observed in biopsies, although these differences did not reach statistical significance (Supplemental Figure 7).

Overall, the small number of differentiated cells recovered by scRNA-seq and the lower transcript abundance limited detection of terminal differentiation genes in organoids. Thus, although IL-13-treated organoids reproduce the overall direction of EoE-associated alterations in terminal differentiation, the day 11 model incompletely captures the magnitude of changes observed in primary tissue. Because transcript abundance may not directly reflect protein expression, we next evaluated expression of the late-differentiation protein, cornulin. Staining showed a preserved organoid organization, with a cornulin-positive differentiated core, which was significantly decreased but not absent in IL-13-treated compared to control organoids (Figure 5B).

Because the number of differentiated cells recovered for scRNA-Seq from esophageal biopsy-derived organoids was low at the day 11 timepoint, we used the EPC2-hTERT organoid model to further characterize organoid developmental kinetics across multiple timpepoints. As previously reported^15^, early differentiation marker, involucrin appears consistently in a small number of cells by day 7 and increases through day 14 whereas cornulin, a marker of later differentiation, became prominent starting at day 9 (Figure 5C).These kinetics suggest timepoints between day 12 and 14 may better capture terminal differentiation and therefore may be optimal for future studies related to terminal differentiation.

### IL-13 Organoids Partially Recapitulate Core Inflammatory Pathways and Cell-trafficking Signals Seen in EoE

We investigated the inflammatory pathways dysregulated in both EoE patients and IL-13-treated organoids. GSEA identified many inflammation-related GO biological processes with shared enrichment, spanning cytokine signaling, innate and adaptive immune processes, and chemotaxis (Figure 6A). Several of these pathways including interferon, TNF, and JAK-STAT signaling have previously been implicated in EoE pathogenesis^40^. The shared leading edge of these pathways included many canonical EoE genes, including *CCL26, POSTN, ALOX15, NTRK1,* and *TNFAIP6* (Figure 6B), indicating that IL-13 alone reproduces much of the core EoE epithelial inflammatory signature.

**Figure 6.**
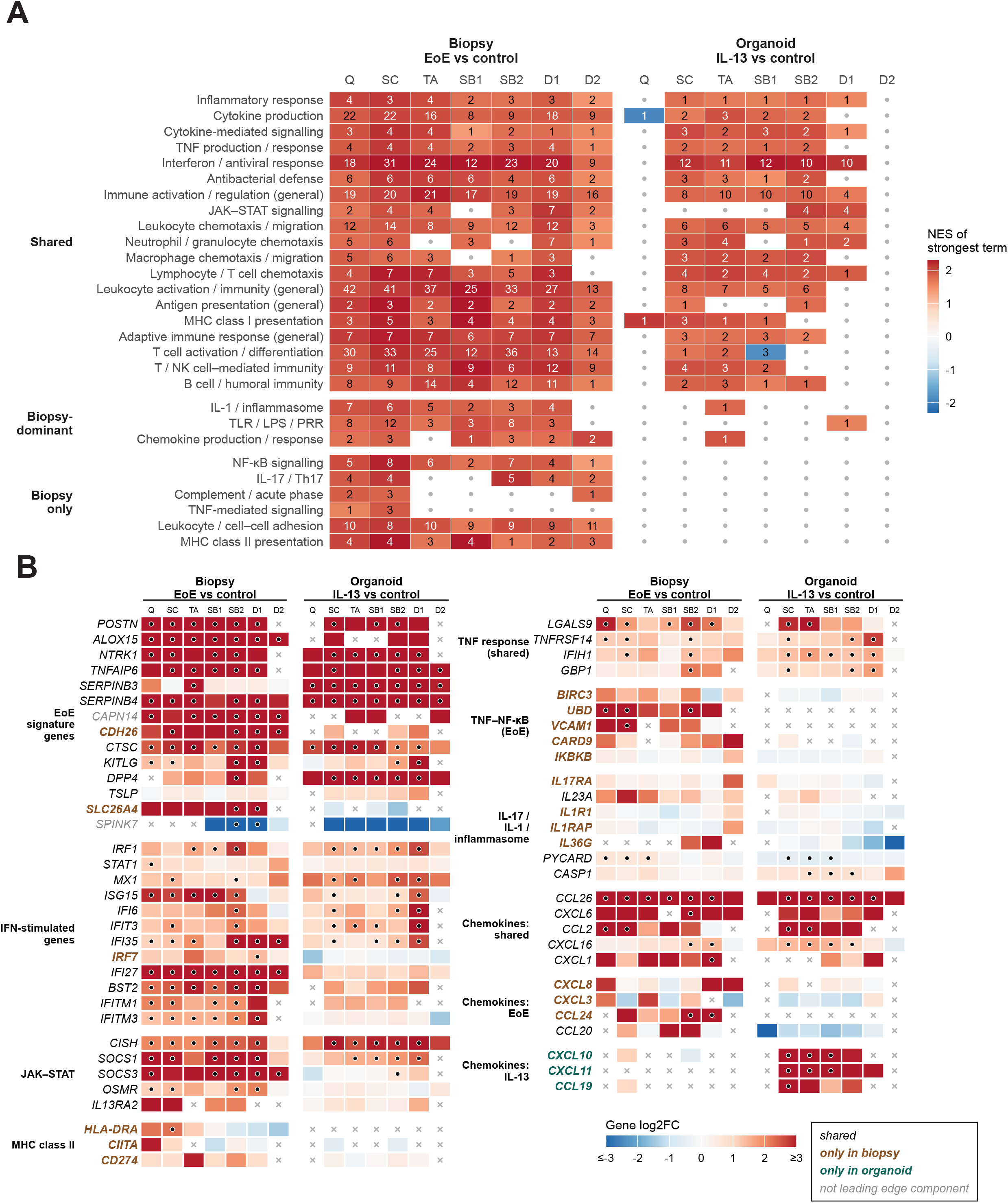
IL-13 organoid models immune activation seen in EoE organoid. A. Inflammation related pathway families significantly enriched in both EoE biopsies and IL-13-treated organoids. Pathways were collapsed into families by theme, and the strongest NES of a pathway-family is shown. The number indicates the number of pathways significant in that family (q <0.05) C. Select leading edge genes of significant pathways. Color indicates log2 fold change. Circle = p adjusted <0.05.

Interferon and JAK-STAT pathways were enriched in both datasets, driven in both by interferon-stimulated genes including *IRF1, STAT1, MX1, ISG15, IFI6, IFIT3,* and *IFI35,* together with IL-13/STAT6 feedback genes *CISH* and *SOCS1.* However, the interferon response was broader in biopsies. *IRF7* only contributed to the leading edge in biopsies, and several ISGs only reached significance in biopsies (*IFI27, BST2, IFITM1/3*). EoE epithelium also showed a STAT3-related response with induction of *SOCS3* and *OSMR,* not seen in IL-13-treated organoids. Similarly, MCH Class II signaling was only enriched in biopsies with *HLA-DRA* and the interferon-induced trans-activator *CIITA* contributing to the leading edge. These findings are consistent with the epithelial interferon signature previously described in EoE^41,42^ and suggest that IL-13 reproduces only part of this response, whereas the full EoE interferon signature likely requires interferon from the tissue environment.

TNF-related pathways also followed a similar pattern. Pathways related to response to TNF were seen in both biopsy and organoid, whereas nuclear factor kappa-light-chain-enhancer of activated B cells (NF-κB) signaling was enriched in only biopsies (Figure 6A, Supplemental Tables 7,8). Additionally, genes contributing to the TNF and interferon pathways leading edges overlapped considerably with roughly 40% of the TNF leading edge genes also contributing to interferon pathways. Several inflammatory pathways were activated in biopsy tissue alone. NF-kB signaling pathways were specific to biopsy, with leading edge including TNF/NF-κB activated genes *BIRC3, UBD,* and *VCAM1* and upstream signaling proteins *CARD9* and *IKBKB* (Figure 6B). IL-17 (leading edge including *IL17RA, IL23A)* and inflammasome/IL-1 signaling pathways were also enriched in biopsy samples.

We next examined epithelial chemokine programs associated with inflammatory cell recruitment in EoE biopsies and IL-13-treated organoids. The canonical EoE-associated eotaxin, eotaxin-3 (*CCL26),* was strongly induced in both EoE biopsy and IL-13 organoid. Eotaxin-3 plays a central role in EoE pathogenesis but does not have a murine ortholog. We examined the expression pattern of *CCL26* in more detail in the organoid model and found that it was broadly induced across all clusters in both biopsies and organoids, with lower expression in the late differentiated compartment (Figure 6B, Supplemental Figure 8). Several other chemokines were also induced in both biopsies and organoids, including *CCL2* and *CXCL16*. In contrast, other chemokines including *CXCL8, CXCL3,* and *CCL24* contributed to the leading edge exclusively in EoE biopsies. Conversely, *CXCL10* and *CXCL11* were highly upregulated in IL-13-treated organoids. This suggests that while IL-13 is sufficient to induce a subset of key chemoattractant signals, the distinct chemokine profile observed in EoE points to additional inflammatory inputs beyond IL-13 alone.

## DISCUSSION

Therapies often fail to resolve the epithelial dysregulation see in EoE, resulting in ongoing symptomatology^5^. Thus, the development of model systems to both understand the pathogenesis of epithelial dysregulation and ultimately to provide a platform for screening of therapeutic targets is critical. Here we build on prior work^15^ by establishing an IL-13-treated 3D organoid system from non-EoE control esophageal biopsies and validating its ability to model changes in the EoE epithelium at single-cell resolution. By defining the cellular composition and transcriptional responses of IL-13-treated organoids, we establish a framework for assessing its fidelity to EoE epithelial biology as well as its potential limitations.

Both control and IL-13-treated patient-derived organoids recapitulated the continuum of differentiation observed in primary esophageal biopsies, preserving a trajectory from basal to differentiated populations. However, the relative cell abundance in the epithelial clusters varied between biopsies and the organoid system, with reduced numbers of quiescent basal and terminally differentiated cells regardless of IL-13 treatment. The scarcity of these populations likely reflects cell culture conditions that favor epithelial growth and self-renewal in organoid culture, resulting in an overrepresentation of proliferative states, These differences highlight an important limitation in the organoid system when modeling the full cellular composition of the native esophageal limitation.

Despite these compositional differences, the IL-13-treated organoid recapitulates the disrupted differentiation dynamics seen in EoE, and both EoE and IL-13-treated organoids show broad dysregulation of epithelial developmental pathways. Several studies have recently suggested that EoE-associated basal cell hyperplasia does not reflect proliferation of basal progenitor cells, but rather an expansion of suprabasal and early differentiated cells retaining basal-like features^10^. Our findings support this model, as *SOX2,* traditionally associated with basal cells, was implicated in gene expression changes across multiple clusters along the differentiation continuum. Additionally, although IL-13 did not change basal cell abundance in the organoids, immunostaining detected an increased proportion of SOX2+ and TP63+ cells, suggesting populations beyond the basal cell compartment contribute to an expanded basal-like phenotype. However, the low number of quiescent basal cells present in the organoid samples limits our ability to assess changes within this population.

IL-13-treated organoids also recapitulated the suppression of terminal differentiation pathways observed in EoE. Late-differentiated cells and their associated transcripts were scarce in day 11 organoids. Protein-based assays may therefore be more sensitive for assessing terminal differentiation in this model and IL-13 treatment was accompanied by a significant reduction in cornulin protein levels at day 11. We hypothesized that extending the culture period to days 12-14 may increase the representation of differentiated cells and improve characterization of late differentiation programs. To evaluate this, we used EPC2-hTERT organoids to evaluate differentiation kinetics across different time points. Compared to patient-derived organoids, the EPC2-hTERT-derived organoids use an immortalized, homogeneous esophageal epithelial population that minimizes inter-individual variability. Although useful for characterizing differentiation dynamics under controlled conditions, this model does not capture the patient-specific or disease-associated heterogeneity represented in patient-derived organoids. We find the proportion of terminally differentiated cells expressing cornulin increases from when it is first expressed on day 9 through day 14. Therefore, harvesting organoids at days 12–14 may increase the yield of terminally differentiated cells, and optimization for a later endpoint should be considered in studies focused on epithelial differentiation.

Lastly, we show that the IL-13-treated organoid recapitulated key inflammatory features of EoE, including increased expression of chemokines implicated in inflammatory cell recruitment during EoE, as well as activation of inflammatory signaling pathways like interferon and JAK/STAT signaling. However, the broader range of inflammatory pathways observed in the epithelium in patient biopsies likely reflects additional inputs from cytokines such as TNF-alpha and TGF-β, as well as other inflammatory cell populations. The interplay between IL-13, interferon signaling, and other inflammatory pathways therefore represents an important direction for future mechanistic studies.

Patient biopsies and esophageal organoids provide complementary approaches to studying EoE-associated epithelial dysfunction. Biopsies capture epithelial changes within the complex inflammatory environment of disease, whereas organoids enable controlled study of epithelial-intrinsic responses to defined stimuli. The ability of IL-13 to recapitulate key features of the EoE phenotype supports the utility of this model for dissecting cytokine-driven mechanisms. At the same time, our data demonstrate differences in epithelial differentiation and inflammatory pathway activation that underscore its limitations in reproducing the full disease environment.

Future refinements in the organoid model, including cytokine co-stimulation, immune or stromal co-culture and optimized differentiation conditions may extend the model’s utility for investigating epithelial-immune interactions. The flexibility and adaptability of the organoid model to these potential adaptations further supports its use in mechanistic and translational studies.

## CONCLUSIONS

In conclusion, our findings establish a framework for evaluating the fidelity of the IL-13-treated organoid model of EoE. This model recapitulates key features of EoE-associated epithelial dysfunction, including dysregulated differentiation, persistence of basal-like transcriptional programs, and activation of inflammatory pathways, while preserving the epithelial differentiation gradient. Although differences in cellular composition and inflammatory signaling were seen between organoids and patient biopsies, the organoid model provides a promising and much needed *in vitro* model to dissect epithelial-intrinsic mechanisms and the contribution of IL-13 to EoE pathogenesis.

## Acknowledgements

We thank the Molecular Pathology and Imaging Core (P30DK050306,RRID: SCR_022420) and the Gastrointestinal Epithelial Modeling Core for technical support. Schematics were created with BioRender.com

## Funding

This work was supported by National Institutes of Health grants 2T32HD043021-19 (A.B), 5KL2TR001879-10 (A.B), K08AI148456 (M.A.R.), R01AI184785 (M.A.R.) and R03DK118310 and R01DK124266-01 (A.B.M.); University of Pennsylvania Transdisciplinary Awards Program in Translational Medicine and Therapeutics (A.B.M.); The Campaign Urging Research for Eosinophilic Diseases (CURED) Foundation (J.B.W), NASPGHAN Foundation Takeda Pharmaceutical Products Inc. Research Innovation Award (J.B.W), Joseph & Dorothy Giddan Child Health Research Award (J.B.W), Ann & Robert H. Lurie Children’s Hospital of Chicago (J.B.W), and The Children’s Hospital of Philadelphia Gastrointestinal Epithelial Modeling Program (T.A.K. and A.B.M.). This work also was supported by P30DK050306: Center for Molecular Studies in Digestive and Liver Diseases.

## COI

The authors disclose the following: Amanda B. Muir has served on the medical advisory boards for Uniquity, Apogee, Esocap, and Sanofi/Regeneron. Joshua B. Wechsler has served on medical advisory boards for Bristol Myers Squibb, Ellodi, Sanofi/Regeneron, and Astrazeneca, and is a speaker for Regeneron/Sanofi, and has received clinical trial funding from Regeneron. Melanie Ruffner has received clinical trial funding from Regeneron. The remaining authors disclose no conflicts.

## Abbreviations

BCH: basal cell hyperplasia
DEGs: differentially expressed genes
DTT: dithiothreitol
EDTA: ethylenediaminetetraacetic acid
EoE: eosinophilic esophagitis
eos/hpf: eosinophils per high powered field
FBS: fetal bovine serum
GO: Gene Ontology
GSEA: gene set enrichment analysis
HBSS: Hank’s Balanced Salt Solution
HEPES: 4-(2-hydroxyethyl)piperazine-1-ethane-sulfonic acid
HPF: high powered field
IBF: impaired barrier function
IL-13: interleukin-13
JAK: Janus kinase
KSFM: keratinocyte serum free medium
KSFMC: KSFM with 1.8 mM Ca2+
PCA: Principal Component Analysis
scRNA-seq: single-cell RNA sequencing
STI: soybean trypsin inhibitor
TA: transit amplifying
TF: transcription factor
Th2: T helper type 2
UMAP: Uniform Manifold Approximation and Projection.

## Data Availability

Esophageal biopsy data will be deposited to dbGAP and accessible upon request or via a data use agreement, organoid data is accessible through gene expression omnibus under GSE227593.

## Ethical Statement

This study was approved by the institutional review board at the Children’s Hospital of Philadelphia (IRB 10-007737) and Lurie Children’s Hospital of Chicago (IRB 2020-3601) and is registered on clinicaltrials.gov (NCT05199532).

Reporting Guidelines: N/A

