## Supplemental Methods for "Systemic Analysis of Patient Derived Esophageal Epithelial Organoid Model Reveals Cell State Heterogeneity and Conservation of Differentiation Hierarchy"

Single-cell RNA-Sequencing (scRNA-seq) Data Processing

Reads were aligned and quantified using 10X Cellranger 3.1.0 and gene expression matrices were generated and loaded into Seurat (V4)^18^. DecontX (v1.2.0) was used to estimate and remove ambient RNA signal from the raw count matrix^19^. Genes expressed in ≤3 cells were excluded. Low quality cells were filtered using criteria: (1) less than 500 or more than 6000 genes per cell and (2) percentage of mitochondrial genes greater than one absolute deviation above the median or greater than 20%, whichever threshold was lower. Doublets were identified and removed using DoubletFinder (v2.0.3), using 10X genomic estimates to set expected double proportion^20^. SCTransform normalization was applied to each sample. Initially, only biopsy and organoid control samples were integrated using Seurat integration workflow (V3/V4) using the top 3000 variable genes. Principal Component Analysis (PCA) was used for initial dimensionality reduction, and the top 45 dimensions of the PCA were used as input for non-linear dimensionality reduction with Uniform Manifold Approximation and Projection (UMAP), cutoff determined by elbow plot. Clustering was performed with a shared nearest neighbor graph by first determining 45 nearest neighbors for each cell.
