## Supplemental Figures for "Systemic Analysis of Patient Derived Esophageal Epithelial Organoid Model Reveals Cell State Heterogeneity and Conservation of Differentiation Hierarchy"

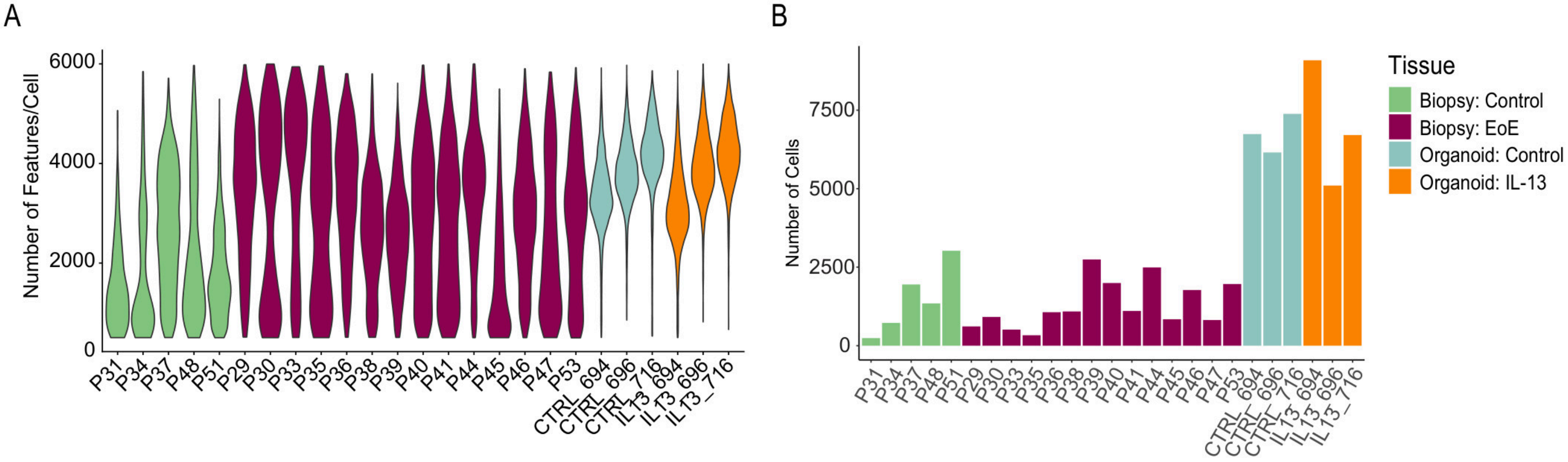


Supplemental Figure 1:Single cell RNA-Seq quality control metrics. A. Number of features per cell by sample B. number of cells per sample. Fill by tissue type.

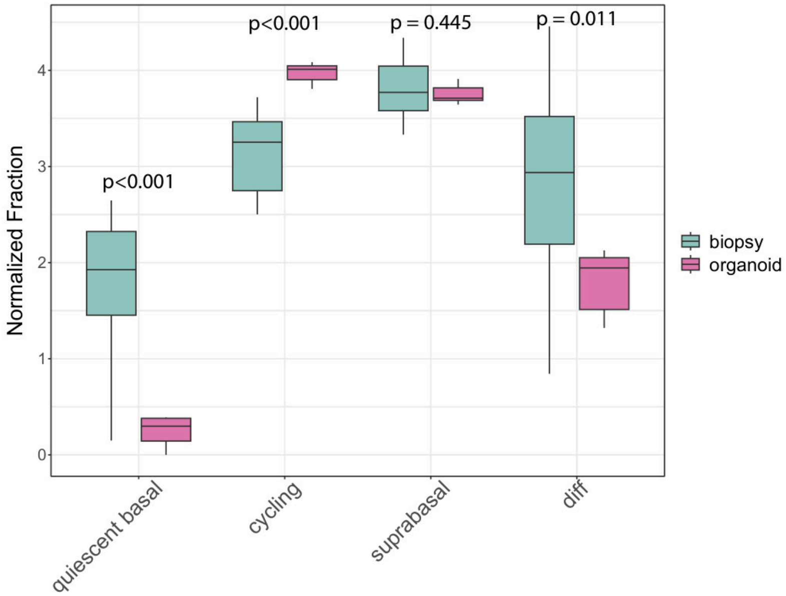


Supplemental Figure 2: Organoids have decreased quiescent basal and differentiated cells as compared to biopsy. Normalized fraction ln(fraction+1) of cell type by tissue type.

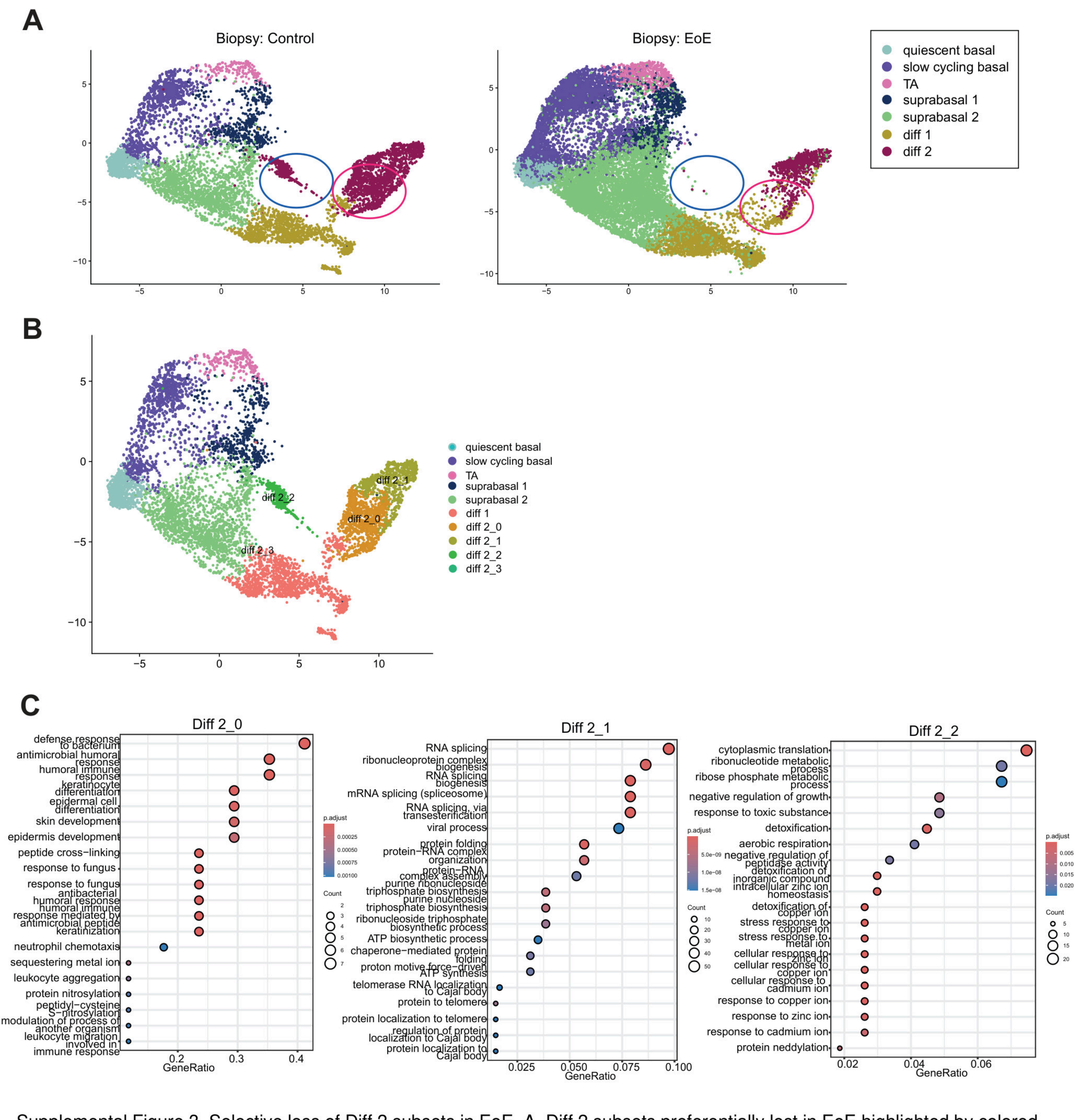


Supplemental Figure 3. Selective loss of Diff 2 subsets in EoE. A. Diff 2 subsets preferentially lost in EoE highlighted by colored circles. B. Subclustering of diff 2 population in control biopsies C. Overrepresentation analysis of differentially expressed genes in GO biologic processes within Diff 2 subclusters

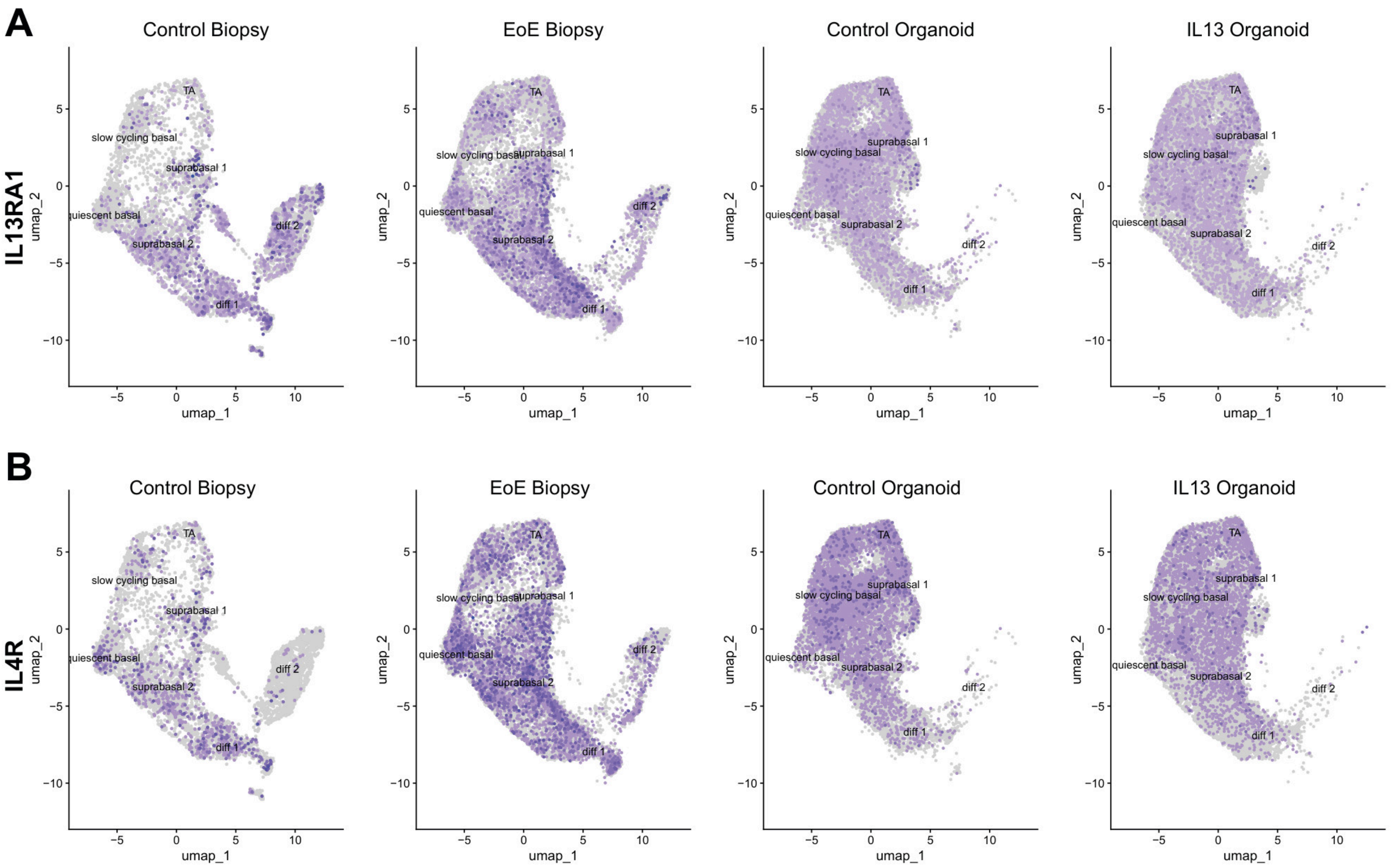


Supplemental Figure 4: IL-13 receptor is broadly expressed across epithelial clusters in biopsy and organoid. Feature plots of IL-13 receptor components (A) IL13RA, and (B) IL4R

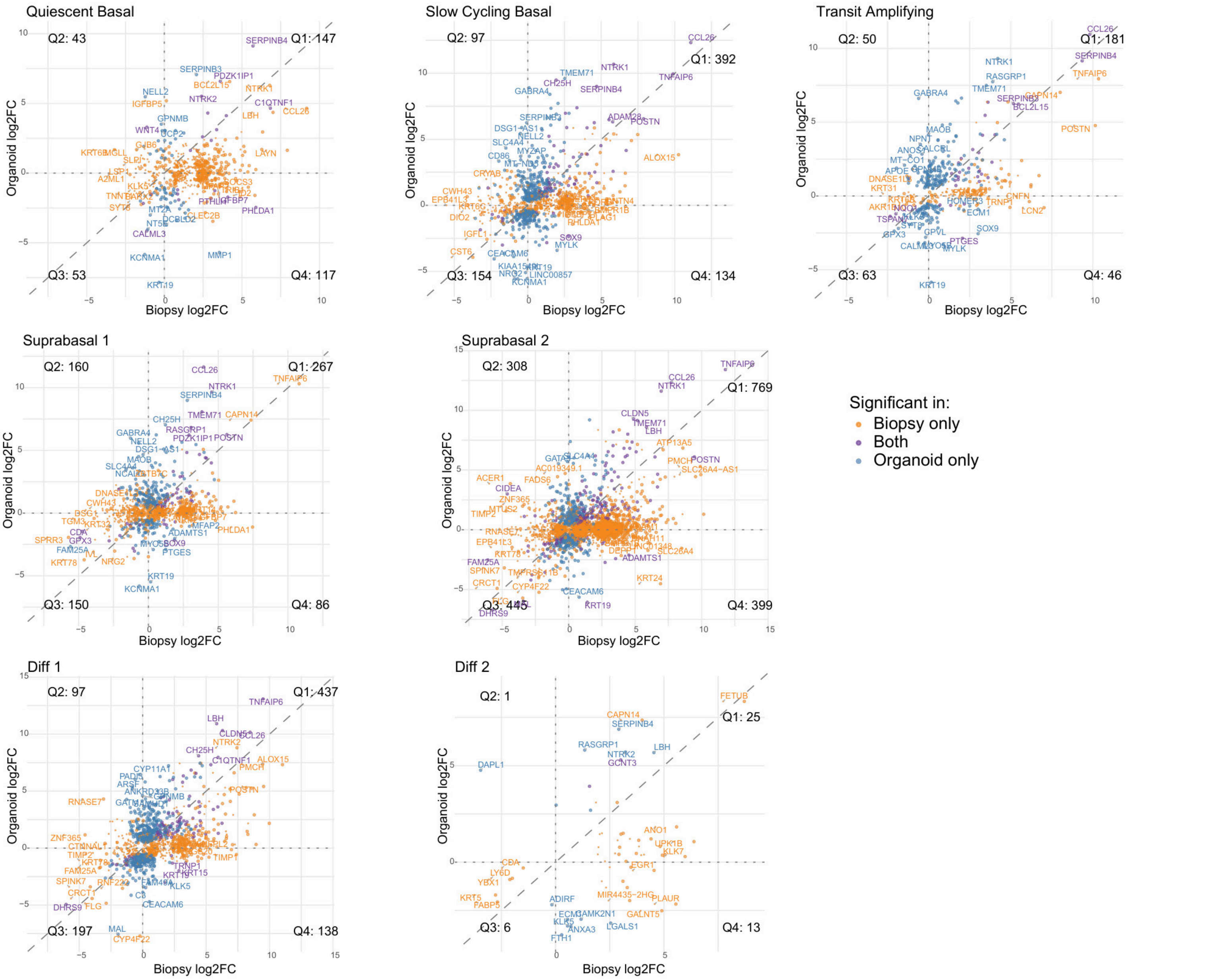


Supplemental Figure 5: Concordance of gene expression changes in EoE and IL-13 treated organoid. Significant genes by cluster, plotted by log2 fold change in biopsy (x axis) versus organoid (y-axis). Color denotes tissue in which a given gene was found significantly differentially expressed (orange – biopsy only, blue – organoid only, purple – both biopsy and organoid). Quadrant numbers denote the number of genes which fall in a given quadrant of the plot.

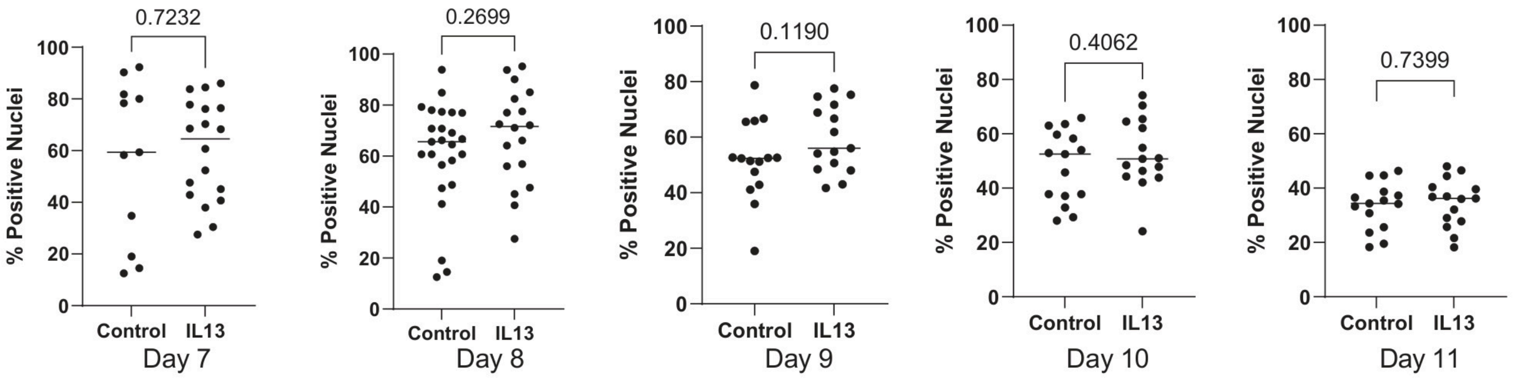


Supplemental Figure 6. Ki67 positivity decreases over time in organoid. Time course of quantification of Ki-67 IHC staining in EPC2 organoids, with or without IL-13 treatment. Mann Whitney U derived p value shown, $p<0.05$ considered significant

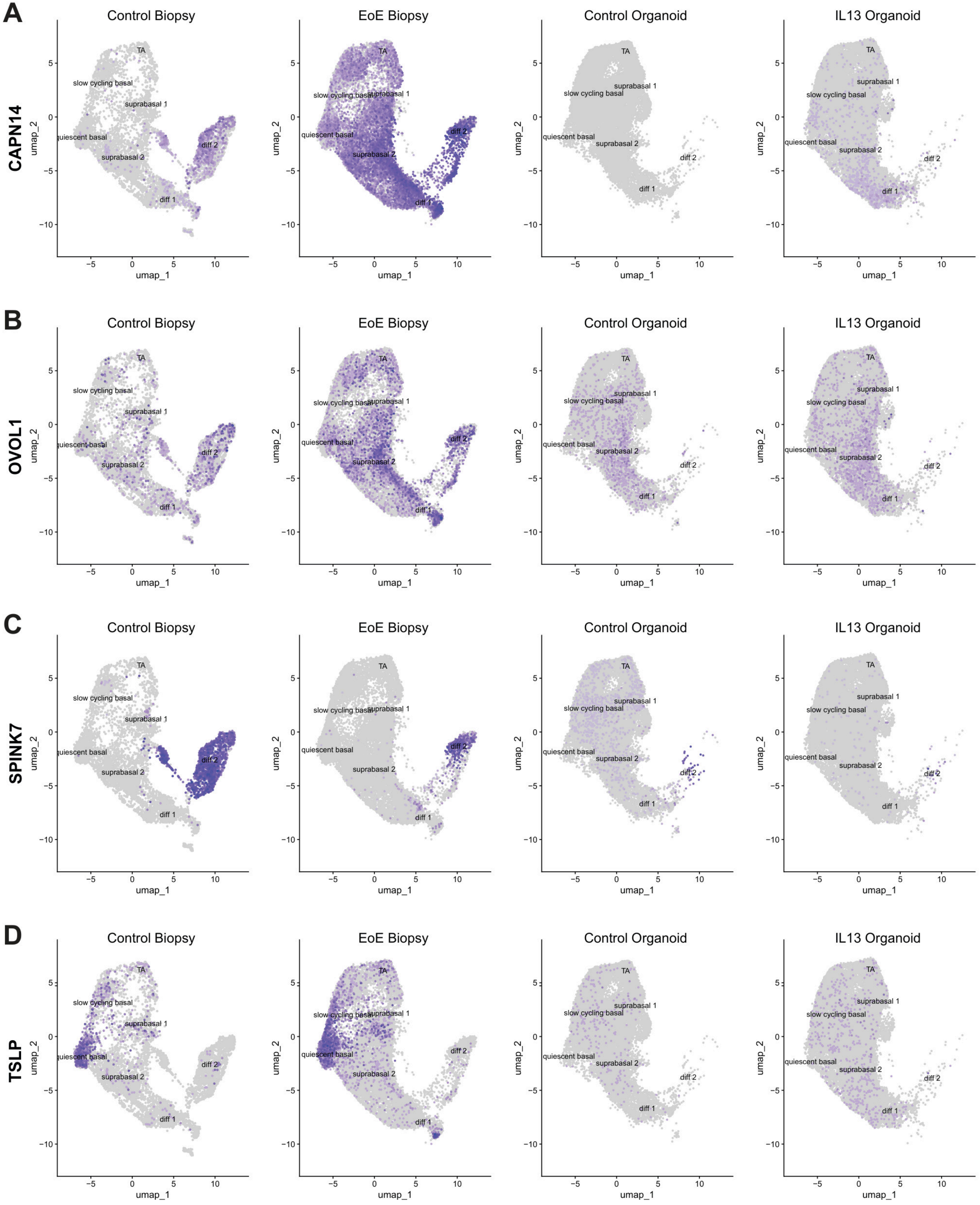


Supplemental Figure 7: OVOL1 pathway components share similar patterns in EoE and IL-13 Organoid. Feature plots of (A) CAPN14 (B) OVOL1 (C) SPINK7 and (D) TSLP expression

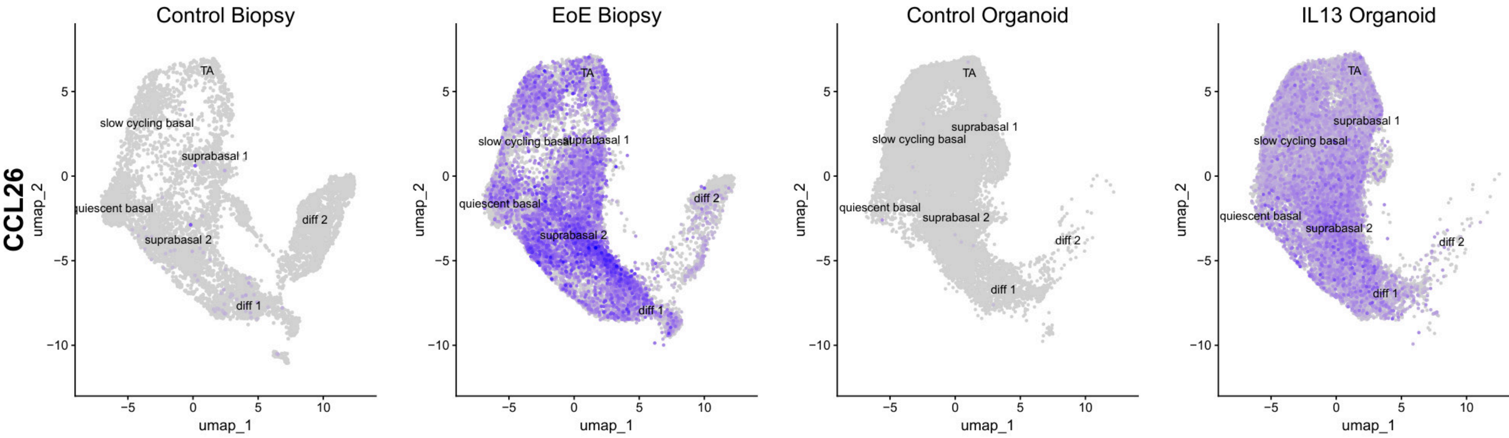


Supplemental Figure 8: CCL26 is broadly induced in EoE and IL-13 treated organoid. Feature plot of CCL26 expression
