## Supplemental Tables for "Systemic Analysis of Patient Derived Esophageal Epithelial Organoid Model Reveals Cell State Heterogeneity and Conservation of Differentiation Hierarchy": SupplementalTable3_PatientCharacteristics.docx

| Active EoE | 100% (14/14) |
| --- | --- |
| Treatment Exposure | 71% (10/14) |
| Dietary Therapy | 64% (9/14) |
| PPI | 57% (8/14) |
| Steroid Exposure | 0% |
| Visual Findings | |
| Exudate | 64% (9/14) |
| Rings | 43% (6/14) |
| Edema | 93% (13/14) |
| Furrow | 86% (12/14) |
| Composite Score Mean [Min, Max] | 4.3 (0-9) |
| Histologic Findings | |
| Maximum Eosinophils/hpf Mean [Min, Max] | 60 (16-100) |
| Basal Zone Hyperplasia  Mild  Severe | 100% (14/14)  50% (7/14)  50% (7/14) |
| Basal Grade  Mild  Mild-Moderate  Moderate  Severe | 14% (2/14)  36% (5/14)  29% (4/14)  21% (3/14) |
| Lamina Propria Fibrosis | 30% (3/10) |

Supplemental Table 2: EoE patient Exposures and Disease Characteristics
